# The evo-MOTiF pipeline and database for studying protein motif evolution in a structural context

**DOI:** 10.1101/2022.08.29.505684

**Authors:** Paul de Boissier, Nathanael Zweig, Bianca H. Habermann

## Abstract

Short linear motifs (SLiMs) in proteins are short functionally independent sequence stretches with a defined function and required for proteins to interact with their environment. Their functional importance makes it interesting to analyse SLiM features further, such as their structural or evolutionary properties, to understand better how SLiMs evolve to shape protein functions.

We developed an automated pipeline to analyse features of SLiMs, called evo-MOTiF. This pipeline takes as input a single protein sequence and its associated SLiM(s) and returns a set of scores associated with SLiM features, including their disorder, as well as their overall, positional and amino acid property conservation. To store and easily mine data from the evo-MOTiF pipeline, we developed the evo-MOTiF database, which currently holds ∼9500 motifs, combining data from ELM, PhosphoSitePlus, as well as from cross-linking mass-spectrometry (XL-MS) experiments. The evo-MOTiF database distinguishes itself further by allowing effortless filtering for SLiMs with specific properties, such as disorder, or conservation in evolution and by providing evolutionary, as well as structural information for SLiMs. Preliminary analysis of SLiM features reveals weak negative correlation between disorder and overall, positional, as well as amino acid property conservation, which is in support of previous observations on smaller datasets.

The evo-MOTiF pipeline and database are freely available at https://gitlab.com/habermann_lab/slims and http://etnadb.ibdm.univ-mrs.fr/index.php, respectively.

## Introduction

Short linear motifs (SLiMs) in proteins are short linear sequence stretches of 3-24 amino acids that are self-sufficient units, required for protein-protein or protein-ligand interaction, with a specific attributed function, such as targeting, degradation, modification, ligand binding and others. As such, they are essential contributors to protein function [1–3].

Functional SLiMs are difficult to predict computationally, due to their shortness and their often-poor conservation. Tools are available for the detection of known SLiMs in protein sequences [4–8], or to detect SLiMs in proteins *de novo* [9–11]. However, predicted SLiMs (*de novo* or not) need to be experimentally verified to provide evidence that they are actually functional [12,13].

The experimental validation of a SLiM for their functional relevance *in vivo* is equally laborious. This explains why the Eukaryotic Linear Motif (ELM) database, the most comprehensive and reliable database of functional and experimentally validated SLiMs, contains, nearly 20 years after its inception, still not more than ∼4000 SLiMs [14]. This number is in relation to the number of proteins known to date and the importance of SLiMs for interaction of proteins with other molecules, probably just the tip of the SLiM iceberg [15,16]. While the ELM database can be seen as the golden standard of SLiMs, methods based on mass-spectrometry have emerged that allow to characterise interacting or modified SLiMs in higher throughput.

With the advent of novel proteomics techniques characterising protein modifications to identify modification sites in proteins, or mass-spec coupled to cross-linking (see e.g. [17–19]) to identify binding sites between two proteins, it has become possible to experimentally find functional SLiMs more easily. While the quality of experimental information on those SLiMs might not meet the high standard of ELM motifs, these identified SLiMs nonetheless contain valuable information on the sequence motif space in proteins.

Davey and colleagues performed a comprehensive analysis of SLiM attributes based on SLiMs stored in the ELM database in 2012 [12]. Using a manually curated sub-set of ELM and by analysing co-crystallized protein structures, they investigated SLiM position, physicochemical properties, evolution and convergent evolution, and the structural context, relating to secondary structure, disorder, as well as overlap with globular domains. Among many interesting observations, two conclusions that are relevant for this study should be pointed out. Observations on the structural context of SLiMs in the protein: SLiMs can be split in those occurring within a conserved domain (partially or completely), while the majority was reported to be outside of such a domain. SLiMs are enriched in disordered regions, as defined by an IUPred [20] disorder score >= 0.4. Both results differ between intracellular and extracellular SLiMs. Observations on SLiM conservation: while motif types like ligand-binding motifs or targeting motifs tend to be conserved over great evolutionary distances, others such as modification sites tend to be relatively poorly conserved, due to their probable redundancy or their simpler nature, allowing them to migrate within a protein. They also confirmed a certain evolutionary plasticity of SLiMs, supported by convergent evolution of certain SLiM types (targeting motifs such as nuclear localisation (NLS) or nuclear export (NES) signals or modification sites), as well as gain and loss of SLiMs in specific lineages. These observations support the hypothesis that SLiMs can evolve rapidly *ex nihilo* or be lost, allowing evolution or loss of protein functions and rewiring the cellular interactome.

*Ex nihilo* evolved SLiMs can have an effect on the function of the protein in which they occur, leading to novel phenotypes [21]. For example, the mutation of a single amino acid in the SHOC2 protein induces the appearance of a N-myristoylation site; The N-myristoyl changes completely the fate of the protein, which then goes to the membrane instead of the nucleus and is implied in rasopathies [22]. Their rather simplistic nature coupled with their enrichment in disordered regions of proteins that typically underlie lower evolutionary pressure, make it tempting to speculate that SLiM evolution could be coupled to the structural context in which they exist and appear. In order to assess this correlation, though, a systematic analysis of existing SLiMs is necessary.

In this study, we set out to develop a straight-forward pipeline and an associated database to store and mine results, called evo-MOTiF, that allows us to extract and correlate features of SLiMs, including their evolution, their secondary structure, their disorder and their surface accessibility. The evo-MOTiF pipeline returns a set of scores describing these attributes and thus enables easy analysis of their correlation to each other. We apply evo-MOTiF to SLiMs from the ELM database [14], to modification sites from PhosphoSitePlus [23], as well as to SLiM regions of XL-MS studies [24] to enlarge the analysed SLiM space. We store results in the publicly available evo-MOTiF database, that contains an exhaustive search mask. This search mask makes analysed SLiMs, together with their scores on disorder, surface accessibility or conservation, easily searchable. Moreover, the evo-MOTiF database offers to view the multiple sequence alignments of the protein families containing SLiMs, as well as 3D-structures of SLiMs alone, as well as in the context of their native protein, if available. The source code of the evo-MOTiF pipeline is freely available at https://gitlab.com/habermann_lab/slims; the evo-MOTiF database is freely accessible at http://etnadb.ibdm.univ-mrs.fr.

## Methods

### The evo-MOTiF pipeline relates protein SLiM evolution to structural context

We set out to develop a pipeline that would help relate the evolution of a protein SLiM to the structural context the SLiM is found in. To this end, we created a pipeline, called evo-MOTiF, that would help us collect this information for a large set of SLiMs without manual interference. We reasoned that we need to collect two crucial pieces of information for this task: first, we needed to identify all orthologs of the protein sequence containing the SLiM. This would help us define the evolutionary history of the motif. Second, we needed to identify the structural state of the SLiM within the protein sequence to help relate SLiM evolution to the structural context of the SLiM. The pipeline is shown in Figure 1.

**Figure 1:**
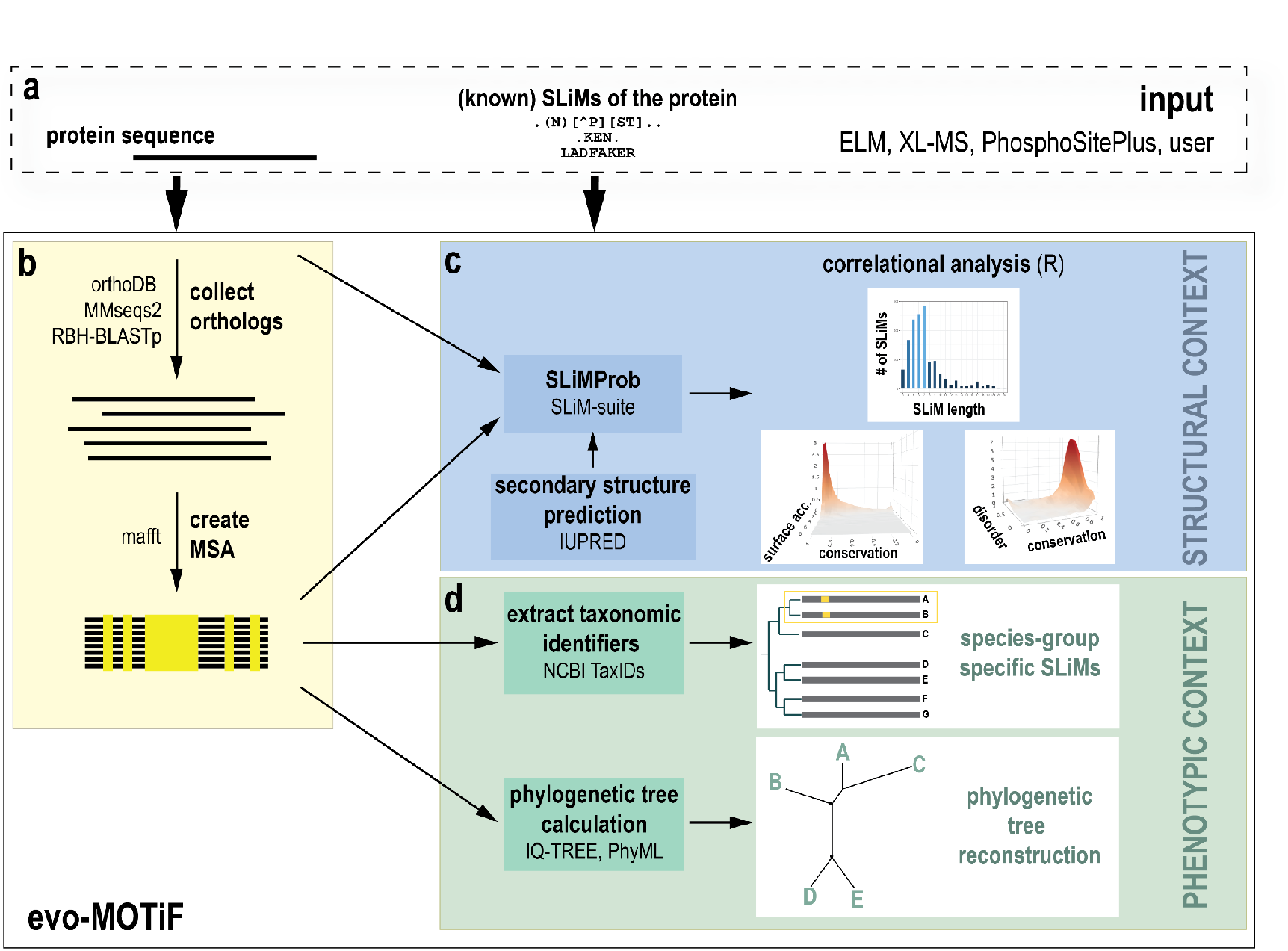
Overview of the evo-MOTiF pipeline. **(a)** evo-MOTiF takes a protein sequence, a list of known SLiMs (either experimentally verified for this protein, or any other SLiM(s) from ELM, PhosphoSitePlus, XL-MS or user-defined) as input. **(b)** First, orthologs of the input protein sequence are searched. This can be done either using data mining of orthoDB, or by running reciprocal MMSeqs2 or BLAST searches. Selected orthologs will be aligned to each other using the MAFFT algorithm, creating a multiple sequence alignment (MSA). **(c)** The MSA, together with the list of SLiMs is submitted to the SLiMProb algorithm. SLiMProb returns a set of scores, including a positional conservation score, a surface accessibility score, as well as a disorder score as defined by the IUPRED algorithm. In addition, we calculate an overall conservation score of the SLiM(s) being analysed. These scores can then be used to perform a set of correlational analyses with the help of a set of R scripts. **(d)** The MSA together with the SLiM information can also be used to deduce evolutionary information for the SLiM. To do so, the Taxomonic Identifier (TaxID from NCBI) is extracted from the MSA, helping to identify taxon-specific SLiMs. An optional phylogenetic tree reconstruction is also offered, using either the IQ-TREE or the PhyML algorithm.

#### evo-MOTiF input

As input, evo-MOTiF requires a protein sequence in fasta-format, as well as a list of SLiMs. For our purpose, this list contained all SLiMs that are associated with the specific input sequence. evo-MOTiF can however also be used to search for SLiMs in an input protein, e.g. by providing the list of known ELM-motifs, or other user-provided SLiMs as input. *Identification of orthologs*. In order to identify orthologs of a SLiM input protein sequence, we encoded three different methods in our pipeline. First, orthologs can be extracted from the OrthoDB database [25]. We chose this database as it is one of the best available orthology databases that has a relatively easy to use API. Second, should the sequence not have any orthologs in OrthoDB, we implemented reciprocal best hit searches, either with the MMSeqs2 algorithm [26], or with BLAST [27]. The best hit, as well as a minimum similarity coverage of 40% was chosen for reciprocal searches. For BLAST, the e-value threshold was set to 0.5. The database for reciprocal searches was reduced to contain only the proteome of the original input species. While MMSeqs2 is much faster than BLAST, BLAST is more sensitive in our hands.

#### Construction of the Multiple Sequence Alignment

The collected orthologs are aligned using the MAFFT program (version 7) using default parameters (FFT-NS-2 method) [28]. The multiple sequence alignment (MSA) is required to further evaluate the evolutionary information for the SLiM under study.

#### SLiM identification

For SLiM identification, we used the SLiMProb algorithm from the SLiM-Suite [4], as it was perfectly suited for this task. It takes the protein of interest together with the SLiM under study (in the form of a regular expression) and the MSA of homologs/orthologs of the protein of interest as input, calculates overall, positional and amino acid property conservation as well as, through the IUPRED program (v1, [20]), a disorder score of the SLiM under study. As output, it produces scores of conservation together with the disorder prediction of the SLiM. Finally, a normalised surface accessibility score is calculated [29]. All scores reported by SLiMProb are also returned by evo-MOTiF and stored in the evo-MOTiF database.

#### Extraction of taxonomic identifiers

In order to make identification of species group- or taxon-specific SLiMs easier, we extract the NCBI TaxID from all orthologs. The TaxID can then be used to relate SLiM presence to the species of the protein.

#### Construction of a phylogenetic tree (optional)

We also offer in the pipeline the construction of an evolutionary tree. This function is optional and should help users to analyse the protein family of the SLiM under study. For phylogenetic tree reconstruction, the user can choose between the PhyML version 3 (with JTT as matrix and 100 iterations for bootstrapping) [30] and the IQ-TREE version 2 algorithm, which produces more accurate phylogenetic trees [31]. This function is optional and can lead to complex trees, depending on the number of orthologs found.

The evo-MOTiF pipeline is freely available for download and local installation from our gitlab repository (https://gitlab.com/habermann_lab/slims).

### Additional necessary processing scripts

We created a script, EXP-MOTiF, that allows the user to choose the original data repository, ELM, PhosphoSitePlus or from XL-MS data, and then prepares data extracted from the respective resources for an evo-MOTiF run. EXP-MOTiF extracts the information about SLiMs from the respective repository and converts UniProt to RefSeq identifiers using the python package MyGene [32]. The RefSeq ID is then used to download the sequence using the Entrez programmatic access [33] and parsed for sequence length. SLiMs are parsed for their corresponding sequences and converted to be readable by SLiMProb. Then, the pipeline is run on each sequence with its specific SLiMs. When SLiM data from ELM are processed, only true positive SLiMs are retained and all other instances of SLiMs found that do not correspond to the true positive ones, are removed. Prior to running SLiMProb, sequences and SLiMs are furthermore pre-processed using the SeqList from the SeqSuite [4].

### Extraction of SLiM structure

To provide structural information for SLiMs, we developed a small pipeline, SLiMs-SCAR (for Short Linear Motifs StruCture extrActoR). It takes evo-MOTiF results in a tsv format as input. Using the submodule PDB from Biopython [34], the corresponding PDB files are downloaded and submitted to our parser, which extracts secondary structure information using DSSP classification [35]. The 2D representation is drawn using Biotite [36]. For displaying the isolated SLiM structure, a PDB file containing the SLiM structure is generated using pdb-tools [37]. 3D visualisation of SLiMs in the evo-MOTiF database is done using JSmol [38].

### SLiM datasets used

We applied the evo-MOTiF pipeline on SLiMs from the ELM database [14], on protein modification sites from PhosphoSitePlus [23], as well as to a set of proteins from cross-linking associated with mass spectroscopy experiments [24].

### Statistical analysis

SLiM data stored in the evo-MOTiF database were further analysed using R. Correlations between SLiM properties were calculated and plots generated with the package corrplot (https://github.com/taiyun/corrplot). Other statistics, such as score distributions and pie charts, were generated using ggplot2 (https://ggplot2.tidyverse.org) and plotly (https://plotly.com/r/).

### The evo-MOTiF database

The evo-MOTiF database was coded using PHP version 7 and HTML version 5. Some plugins are implemented using JavaScript version 1.7. The entries of the database are stored using MySQL version 5. Data from the MySQL database are extracted through PHP and JavaScript. We used the plug-in DataTables (https://www.datatables.net/) to design the main page of the database in which every motif is stored and can be searched. For each motif, a page is automatically generated, extracting information from the protein XML file from UniProt and the MySQL database. JSmol is used to display PDB structures in 3D, allowing users to change the appearance of the visualised protein- and SLiM structures.

The structure of the evo-MOTiF database is shown and further discussed in Supplementary Figure S1.

## Results

### Description of SLiM scores calculated by the evo-MOTiF pipeline

Several scores are calculated and/or returned by the evo-MOTiF pipeline which are described in the section below.

#### Disorder

The disorder score describes the structural state of the motif. It is likewise returned by SLiMProb and is calculated using the IUPRED algorithm. It ranges from 0 (structured) to 1 (disordered).

#### Overall conservation

The overall conservation score is reported by the SLiMProb algorithm and gives the proportion of sequences, in which the SLiM under study is conserved. It ranges from 0 (overall not conserved) to 1 (overall conserved).

#### Positional conservation

The positional conservation score is reported by the SLiMProb algorithm and describes the conservation of each residue of the SLiM on the basis of the MSA given to the algorithm. The score ranges from 0 (not conserved) to 1 (conserved).

#### Amino acid property conservation

The amino acid property conservation score is reported by SLiMProb and gives a conservation score based on the properties of amino acids for the SLiM. The default amino acid property matrix suggested by SLiMProb is used. It ranges from 0 (not conserved) to 1 (conserved).

#### Surface accessibility

The surface accessibility gives the probable accessibility of the SLiM with respect to its immediate neighbourhood of amino acids. It is calculated by SLiMProb. It ranges from 0 (buried) to more than 9 (accessible).

#### Probability

The probability is extracted from the ELM database and gives the probability, a specific SLiM exists in the protein. This score is only available for ELM motifs.

All scores calculated by the evo-MOTiF pipeline are searchable in the evo-MOTiF database.

### The evo-MOTiF database

We developed the evo-MOTiF database, accessible at http://etnadb.ibdm.univ-mrs.fr, to allow an easy mining of protein SLiM evolution in a structural context. A schematic representation of the evo-MOTiF database is shown in Figure 2.

**Figure 2:**
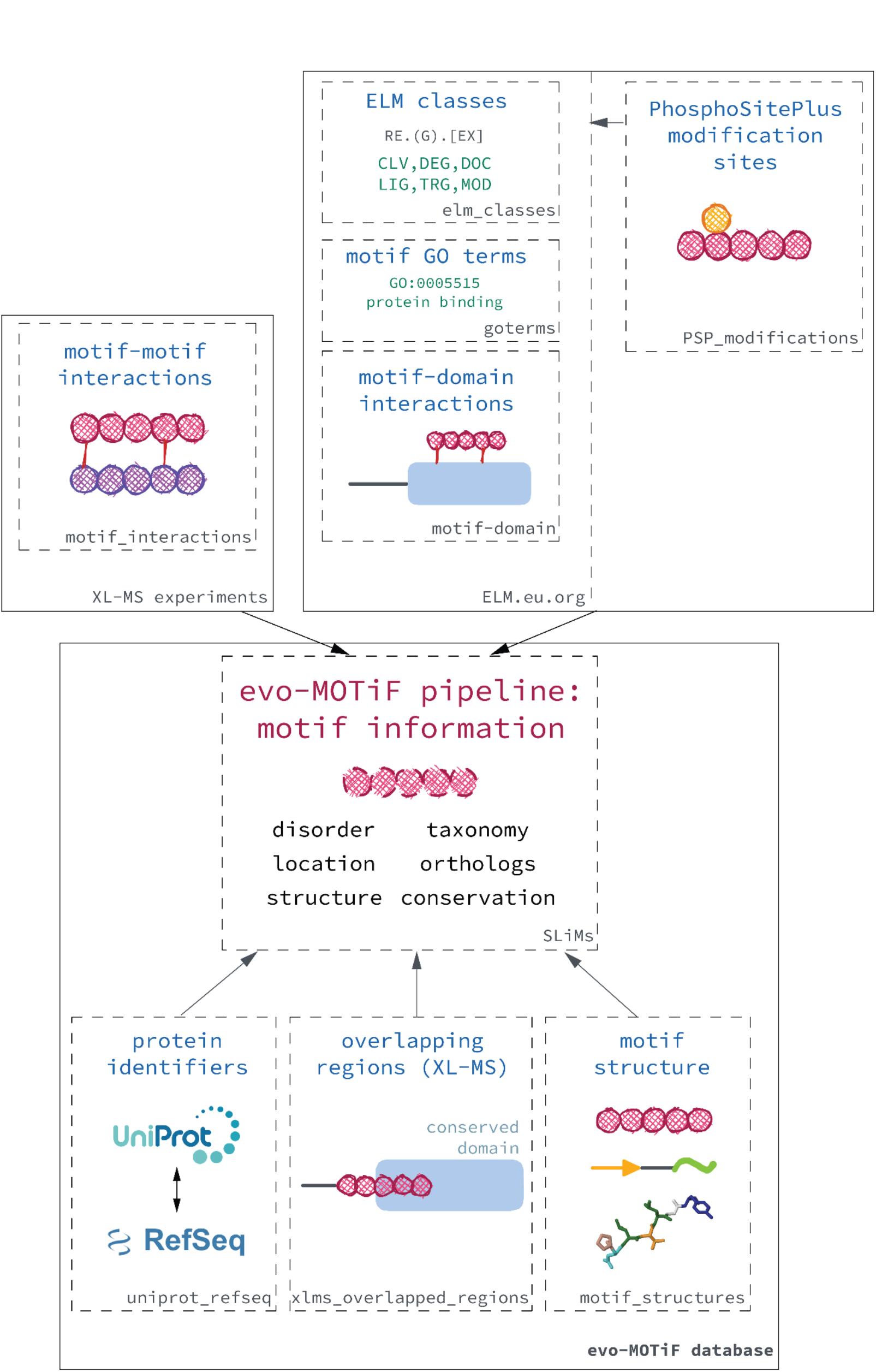
Schematic representation of the evo-MOTiF database. SLiMs are extracted from ELM, PhosphoSitePlus, as well as from XL-MS experiments. In case of ELM motifs, ELM motif classes, GO annotation, as well as motif-domain interaction information are extracted. PhosphoSitePlus motifs are conformed to ELM motifs, and are assigned the new motif class PSP_modification. Motif-motif interactions are extracted from XL-MS data. These experimentally verified SLiM(s) are together with their respective protein sequence submitted to the evo-MOTiF pipeline, which calculates scores such as positional and overall conservation of the SLiM, provides taxonomic information, calculates surface accessibility, and disorder scores, as well as extracts the structure of the SLiM, if a three-dimensional structure is available. Next to these data, the Uniprot and RefSeq Identifiers are stored, overlapping conserved domains to XL-MS motifs are extracted and structural information (from pdb) are given.

We collected protein SLiMs from the ELM database [14], PhosphoSitePlus [23], as well as from XL-MS experiments [24]. Each SLiM, together with the full-length protein sequence, were submitted to the evo-MOTIF pipeline to find orthologs, create the MSA, refind and score the SLiM in the protein sequence and its orthologs, and collect information on the overall evolution (i.e. the presence of the SLiM in the orthologs), the context of the secondary structure (helical, beta-strand, unstructured/disordered), as well as the taxonomic identifier of the protein and all orthologs. In addition to these core scores, the structure of the protein and its SLiM were subtracted from the protein data bank (PDB) database and is accessible at a pop-up page to view from the database. Additional information was extracted from the different resources and were stored: the interacting domain (ELM) or motif (XL-MS data); ELM motif classes (ELM), as well as Gene Ontology (GO) information (ELM).

The main database page is shown in Figure 3. In the database table, each motif is shown, with identifiers, origin, disorder and conservation scores, surface accessibility, as well as the taxonomic identifier of the protein the SLiM originates from. Each column can be individually searched, to filter for SLiMs with a specific feature, using SLiM scores such as disorder, conservation, etc. or the taxID / species. A general search field at the top of the table also allows to perform free-text searches, applying to all fields of the database. The individual search fields for the SLiM scores makes it exceptionally easy to filter for SLiMs that have certain amino acid property-, positional- or overall-conservation, disorder, or surface accessibility scores. This filtering function helps to quickly find SLiMs that are found in highly disordered or ordered regions of the protein, or those that have high-, or low conservation.

**Figure 3:**
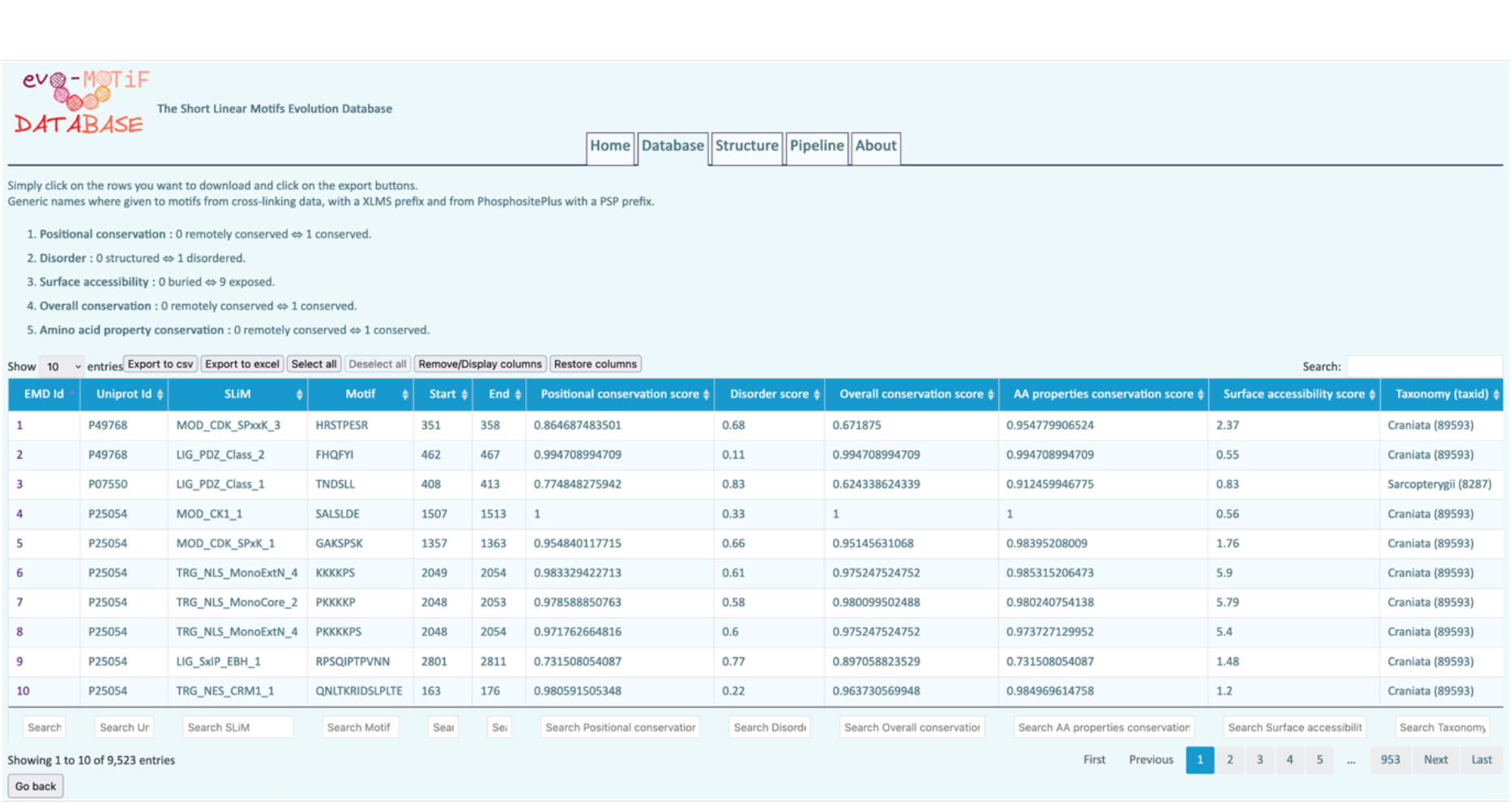
Main page of the evo-MOTiF database. SLiMs are shown in tabular format, with information on the SLiM itself, the position in the sequence, several scores provided by the evo-MOTiF pipeline, as well as the taxonomic identifier. Each column is searchable, which makes the evo-MOTiF database a powerful tool to search for SLiMs based on their evolution, or their structural properties (ordered, disordered structures). A free-text search at the top of the page also allows searching all fields simultaneously.

The database page for an individual SLiM (Supplementary Figure S2) contains a short information section on the protein, and information on the SLiM, including the regular expression, sequence and a description of the associated function, with the link to the original source (ELM or PhosphoSitePlus). This section also contains information on the evolution of the protein, with the number of orthologs, as well as a link to view the multiple sequence alignment of the protein family (Figure 4 b). This is followed by the scores calculated by evo-MOTiF (Figure 4 a) and the interacting domain (in case of ELM) or SLiM (in case of XL-MS motifs). At the next section, the GO annotation of the SLiM is provided. At the bottom of the web-page (Supplementary Figure S2), all SLiMs are shown for the protein, schematically, as well as in a table. The structure of the selected SLiM is given, as is a link to the structure page of the SLiM, which is shown in Figure 5.

**Figure 4:**
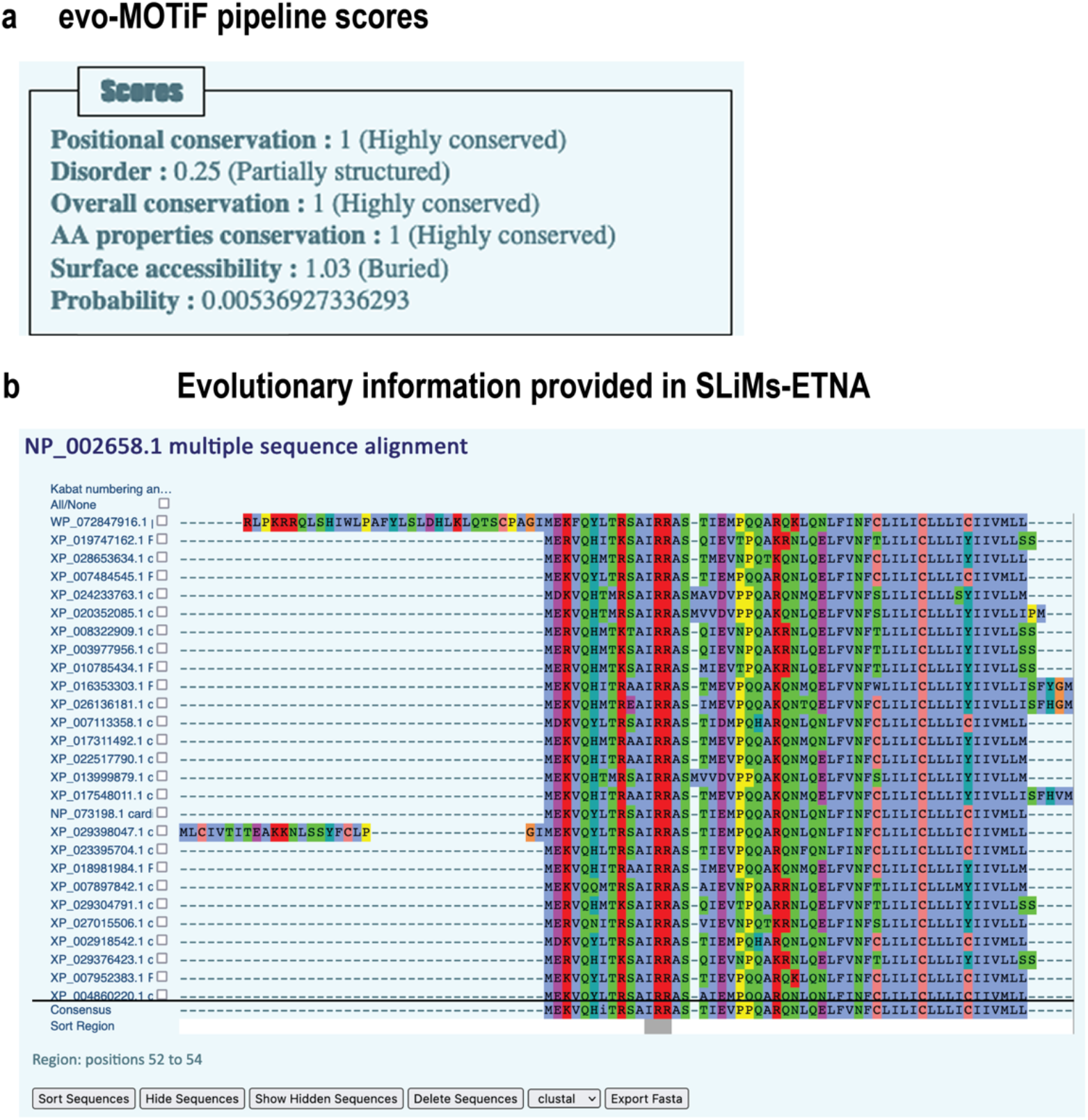
Representation of the Multiple Sequence Alignment of an evo-MOTiF entry. **(a)** The scores returned for a SLiM in the evo-MOTiF database. Next to amino acid property, positional and overall conservation, a disorder score, the surface accessibility, and - for ELM motifs - also a probability is given. **(b)** Multiple sequence alignment of an evo-MOTiF entry. The alignment is displayed using the JSAV plugin, offering many functionalities, such as sorting according to a selected sequence stretch, hiding and deleting sequences, colouring of the alignment using different colouring schemes, or exporting the alignment as fasta. For Figure 3, the SLiM entry 751 (P26678, PPLA_HUMAN - Cardiac phospholamban) is used, which contains a TRG_ER_diArg_1 motif.

**Figure 5:**
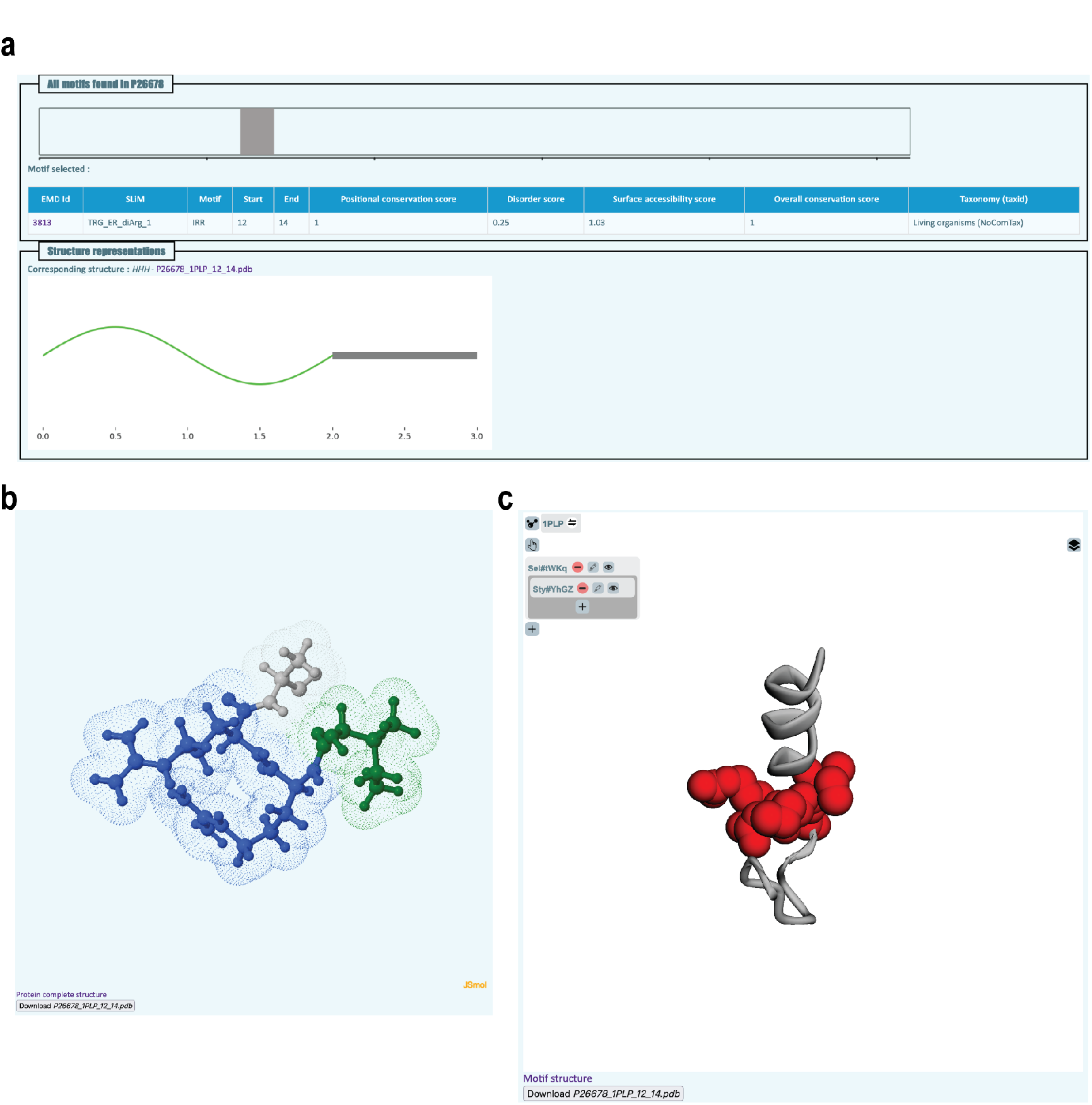
Structural representation of a SLiM entry in the evo-MOTiF database. (a) The sequence is represented as a bar, with the motif highlighted in grey. The nature of the motif is shown at the bottom (in this case, helical and unstructured). (b) 3D structure of the motif itself, displayed with the JSmol plugin. A ball-and-stick representation is chosen, coloured according to amino acid. In addition, a dotted surface is displayed. (c) To show the entire protein structure, with the motif highlighted in red, the 3Dmol plugin is used. The structure is shown in cartoon mode in grey, with the SLiM itself in space fill in red. The user can toggle between the display of the motif alone, or the motif in the context of the entire protein structure. For Figure 4, the SLiM entry 751 (P26678, PPLA_HUMAN - Cardiac phospholamban) is used, which contains a TRG_ER_diArg_1 motif.

One of the main features of the evo-MOTiF database is the direct availability of evolutionary as well as structural information on the protein and the SLiM. Figure 4 shows the MSA of the protein family, displayed via the JSAV plugin [39]. Figure 5 shows the structure of the SLiM, displayed using the JavaScript plugin JSmol [38] (http://www.jmol.org/), accessible via the main page, as well as the structure of the entire protein, in which the SLiM is highlighted in red, through the JavaScript plugin 3Dmol [40].

### Statistical information on the evo-MOTiF database

The evo-MOTiF database at the time of writing holds 9523 SLiMs, with 4733 from XL-MS data, 1982 from ELM and 2808 from PhosphoSitePlus (Figure 6 a).

**Figure 6:**
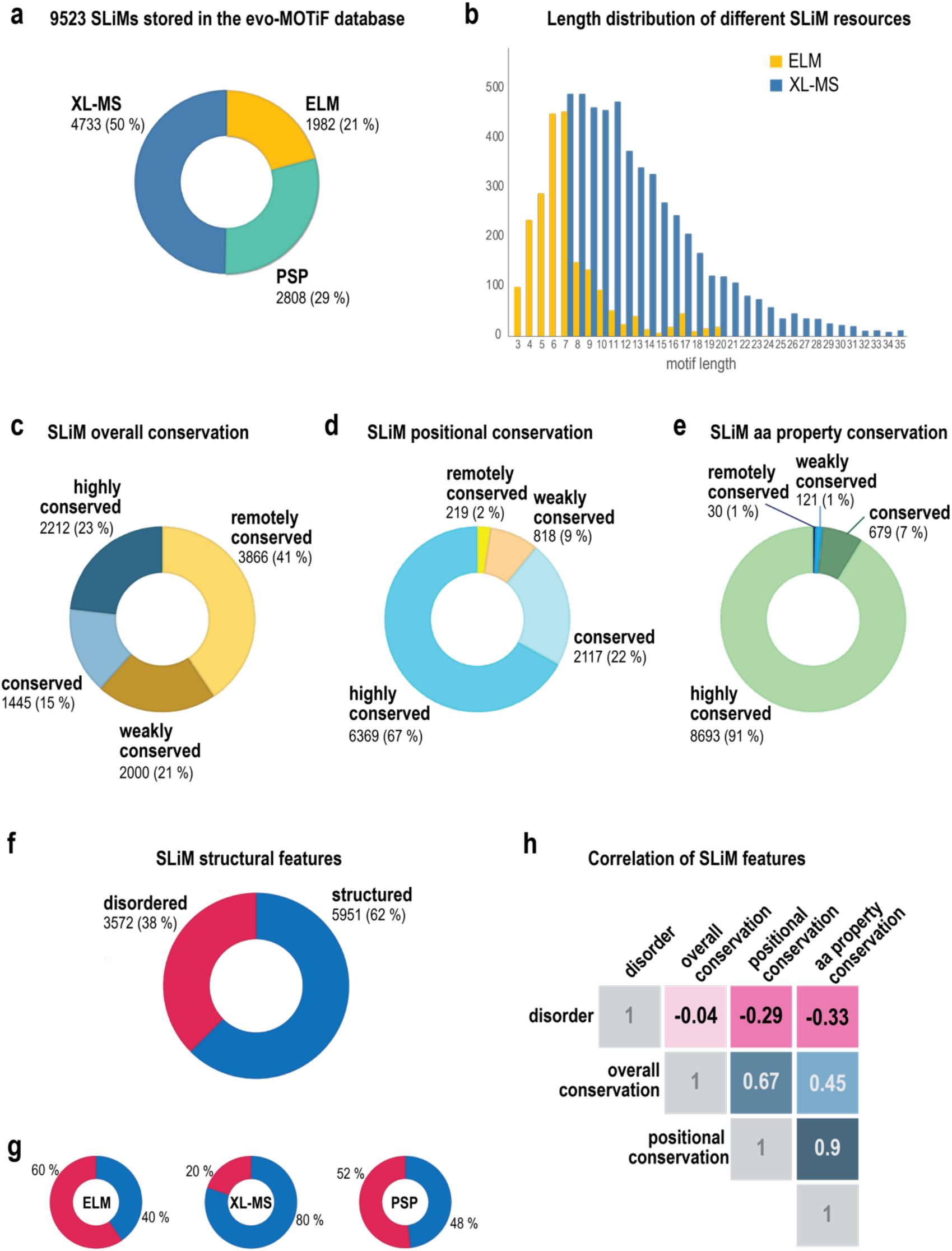
Preliminary statistical and correlational data extracted from processed SLiMs of the evo-MOTiF database. **(a)** Number of SLiMs already processed and available via the evo-MOTiF database. **(b)** Length distribution of SLiMs from ELM and XL-MS data. ELM SLiMs have an average length of 7, SLiMs from XL-MS have an average length of 13 residues. XL-MS motifs are also generally longer and are not shorter than 7 amino acids. **(c)** Overall conservation of SLiMs in the evo-MOTiF database. Only 23% of motifs can be considered highly conserved and more than 43% of SLiMs fall into the category ‘remotely conserved’, with an overall conservation score <0.25. **(d)** Positional conservation of SLiMs available in the evo-MOTiF database. 67% of SLiMs have high positional conservation (positional conservation score > 0.75), 2 % of motifs have low or remote positional conservation. **(e)** Amino acid property scores of analysed SLiMs. The properties of amino acids are highly conserved in more than 90% of motifs. **(f)** Structural features of SLiMs available in the evo-MOTiF database. Over all three resources, the majority of SLiMs is structured (62%). **(g)** Structural features of the individual resources. More SLiMs from ELM are disordered (60%), while only 20% of SLiMs from the XL-MS experimental data are considered disordered. PSP motifs have a nearly equal distribution between disordered and structured motifs. **(h)** Correlation analysis of SLiM structural and evolutionary scores. A strong positive correlation can be seen between the three conservation scores. Disorder is very weakly negatively correlated with overall conservation and shows moderate negative correlation with positional, as well as amino acid property conservation. All correlation values are significant (p-value < 2E-04).

SLiMs from ELM and XL-MS show marked size differences: The distribution of SLiM lengths is different, with ELM SLiMs peaking at length 6 and 7 and a mean motif size of 7 amino acids, while XL-MS motifs are not shorter than 7, and a mean motif size of 13 amino acids (Figure 6 b). XL-MS SLiMs can also be as long as 35 amino acids. PhosphoSitePlus motifs, unless located in N- or C-terminus of the protein, have a size of 15, due to technical reasons of mass-spec motif discovery (data not shown).

Many of SLiMs (41%) are overall only remotely conserved, with an overall conservation score of <0.25. Only 23% of motifs are highly conserved (Figure 6 c). For the vast majority of SLiMs, the average positional conservation is high (67%), 22% can still be considered conserved (positional conservation score > 0.5), and 11% have low or remote positional conservation (Figure 6 d). With over 90% of SLiMs having an amino acid property score >0.75, the conservation of amino acid properties of motifs is generally very high (Figure 6 e). There are striking differences between the 3 resources (ELM, XL-MS and PhosphoSitePlus), especially concerning the overall conservation. The majority of ELM motifs are in fact, highly conserved (66%), while only 13%, and 10% of motifs from XL-MS and PhosphoSitePlus are highly conserved, respectively (Supplementary Figure S3 a). SLiM positional conservation is more similar between resources, especially ELM and XL-MS motifs (77% in both are highly conserved on positional level). Only PhosphoSitePlus motifs have a lower % of high conservation on positional level (43%) and more motifs that have a positional conservation score below 0.5 (Supplementary Figure S3 b). Finally, amino acid properties of SLiMs are well conserved across all three resources (Supplementary Figure S3 c).

Interestingly, the majority of SLiMs in our database are defined as structured (with an IUPRED score <0.5 [41], 62%, Figure 6 f). However, SLiM disorder peaks around 0.5 and most SLiMs are found close to this cut-off, being either partially structured, or partially disordered (see Supplementary Figure S4). We found this result surprising, because of the described preference of SLiMs for disordered regions [12]. We therefore took a closer look at the disorder distributions of the three different resources (Figure 6 g). The majority (60%) of ELM motifs are indeed found in disordered regions, while 80% of SLiMs from the XL-MS study analysed are found in structured regions, indicating a bias in the resource, which might be due XL-MS motifs being slightly longer. PhosphoSitePlus SLiMs show an even distribution between structured and disordered regions. It should be noted here that in [12], following recommendations of the developers of SLiMSearch [11] and SLiMPrints [10] for motif finding in disordered regions, an IUPRED cut-off of 0.4 was chosen to distinguish between a sequence being structured or disordered. Using this cut-off changes the interpretation of our data and flips the balance of SLiM structural properties, with 52% of all evo-MOTiF database motifs being disordered and 48% being structured.

We finally looked at the correlation between structural and evolutionary features of SLiMs (Figure 6 h). There is high positive correlation between the conservation scores, as is expected. Disorder is negatively correlated with all three conservation scores, whereby negative correlation with overall conservation is very low (0.04), however statistically significant (Supplementary Table S1 i). This finding is in agreement with previous hypotheses and observations that structural disorder may be related to SLiM evolution [12,21]. Correlation plots for the individual resources are shown and further discussed in Supplementary Figure S5.

All statistical analysis of SLiM features done is also provided for further mining in the evo-MOTiF database (http://etnadb.ibdm.univ-mrs.fr/statistics.html).

## Discussion

We present here a new pipeline and resource for short linear motifs in proteins, the evo-MOTiF pipeline and database. The evo-MOTiF pipeline was designed to ease analysis of short linear motifs and allow to easily find and analyse SLiMs with different structural, as well as evolutionary features. We were and are analysing SLiMs from ELM, XL-MS experiments, as well as from PhosphoSitePlus with the evo-MOTiF pipeline. To effortlessly mine those data, we store the results in the evo-MOTiF database, which has an extensive search interface and allows searching and filtering for SLiMs with certain properties, by filtering for scores of their disorder, and their overall, positional, or amino acid property conservation.

We wanted to systematically analyse SLiMs for their properties to get a better picture on their evolutionary or structural features. We designed our pipeline such that it would run automatically, given a (user) input of a sequence and its associated motifs, perform all the steps required and return scores for several SLiM properties relating to their structure and evolution. Among the challenges in this workflow are the choices for reliably finding and correctly aligning orthologs. We chose to offer three different workflows for finding orthologs: reciprocal BLAST-searches, reciprocal MMSeqs2-searches and mining orthologs from the orthoDB database. While BLAST is the most sensitive method, it is also the computationally most expensive one. We therefore chose to run our reciprocal homology searches using MMSeqs2. We aligned identified orthologs using the MAFFT algorithm, which is reliable and fast. Another choice that had to be made was to select the algorithm to search for known SLiMs. We chose the SLiMProb algorithm, as it already returns some of the scores we were interested in. We could effectively use the three conservation scores, as well as the disorder score, while we chose to not further consider the surface accessibility returned by SLiMProb. According to the resulting surface accessibility scores, more than 90% of SLiMs were buried, indicating that this measure has little use for further distinguishing SLiM properties. This score nonetheless remains to be accessible in the evo-MOTiF database.

The evo-MOTiF pipeline is computationally expensive, which is why only a portion of available motifs have so far been processed. While we make the pipeline available to others, it should be noted that a high-performance computer is required to run it. Improvements could be made in making the pipeline parallelized and suitable for a high-performance computer cluster.

We designed the evo-MOTiF database as a complementary resource to other motif databases, such as foremost ELM, as well as the PhosphoSitePlus database. Each motif in the evo-MOTiF database is linked to its original source, so its origin should be clear to evo-MOTiF database users. We wanted to create a database that contains only experimentally identified SLiMs, but not be as exclusive as ELM. All SLiMs found in evo-MOTiF have experimental proof, while only motifs originating from ELM meet the ‘golden standard’ criterion of a *bona fide* protein motif. PhosphoSitePlus motifs are less deeply curated and XL-MS motifs are by definition originating from primary experimental data without further manual curation or verification of the motifs. The evo-MOTiF database is in fact the first motif resource that stores XL-MS data in this form. P-values, providing a certainty to the functionality of the motif, are extracted from, and thus only given for ELM motifs. As we do not further curate SLiMs from the other resources, the high standard of ELM is also only available for ELM motifs stored in our database.

Next to a broader range of motifs stored, the evo-MOTiF database distinguishes itself from other motif databases by further functional features: only our database has the possibility to effortlessly search for individual motifs based on their properties, such as structural disorder, or conservation attributes. It is for instance possible to search for overall remotely conserved - or *ex nihilo* born - motifs by simply filtering for motifs with a low overall (or positional) evolutionary score. The evo-MOTiF database provides identified orthologs of a motif-carrying protein and allows users to download, or view the multiple sequence alignment of all orthologs. The evo-MOTiF database provides secondary structure predictions, as well as disorder predictions of SLiMs and allows users to filter for structural SLiM attributes. Finally, the evo-MOTiF database provides and displays structural information, if available, of a SLiM alone, as well as in the context of the entire protein structure. It is therefore possible to easily visualise the position of the SLiM in the overall 3D structure, which might aid in further analysis of the SLiM and its function.

Our preliminary statistical analysis of SLiMs stored in the evo-MOTiF database revealed that some features are specific to the source of the SLiMs: ELM, PhosphoSitePlus or XL-MS data. For instance, we found a difference in length distribution depending on the source. Motifs found with XL-MS tend to be longer than ELM motifs. This length difference most likely impacts some other difference in motif features we have observed. More specifically, the fact that XL-MS motifs are generally more structured than ELM motifs might be due to the fact that they are longer in sequence. Most of the SLiMs also have a IUPRED score close to 0.5. Currently, a cut-off of 0.4 is advised from the developers of SLiMSearch and SLiMPrints to find SLiMs in *disordered* regions, while the fact seems to be that most SLiMs are in partially structured or partially disordered regions. We specifically chose the generally recommended cut-off of IUPRED (version 1) of 0.5, as the method predicts disorder not only of a motif, but returns a disorder score over a region of a 21 amino acid window and in that sense takes more into account than just the motif sequence. We want to generally raise the question, whether IUPRED 1.0 is suitable for predicting SLiM structural features, such as disorder, or whether a more motif-specific algorithm should be considered, such as IUPRED2 or 3 that allow to predict short disordered regions. Furthermore, with the current data, it raises the question, whether any hard cut-off is useful in further analysing SLiM structural features.

Finally, we observed a negative correlation of overall, positional as well as amino acid property conservation with SLiM structural disorder across all three resources, further supporting the hypothesis that faster evolution rates in disordered regions might assist SLiM evolution. This is not unexpected and has been observed and discussed previously, though based on a set of different SLiM data and using different workflows to calculate SLiM attributes [12].

