## Supplementary data for "The evo-MOTiF pipeline and database for studying protein motif evolution in a structural context"

#### Supplementary Information

Supplementary Figures S1 - S5

#### Supplementary Figure S1

### evo-MOTiF database structure

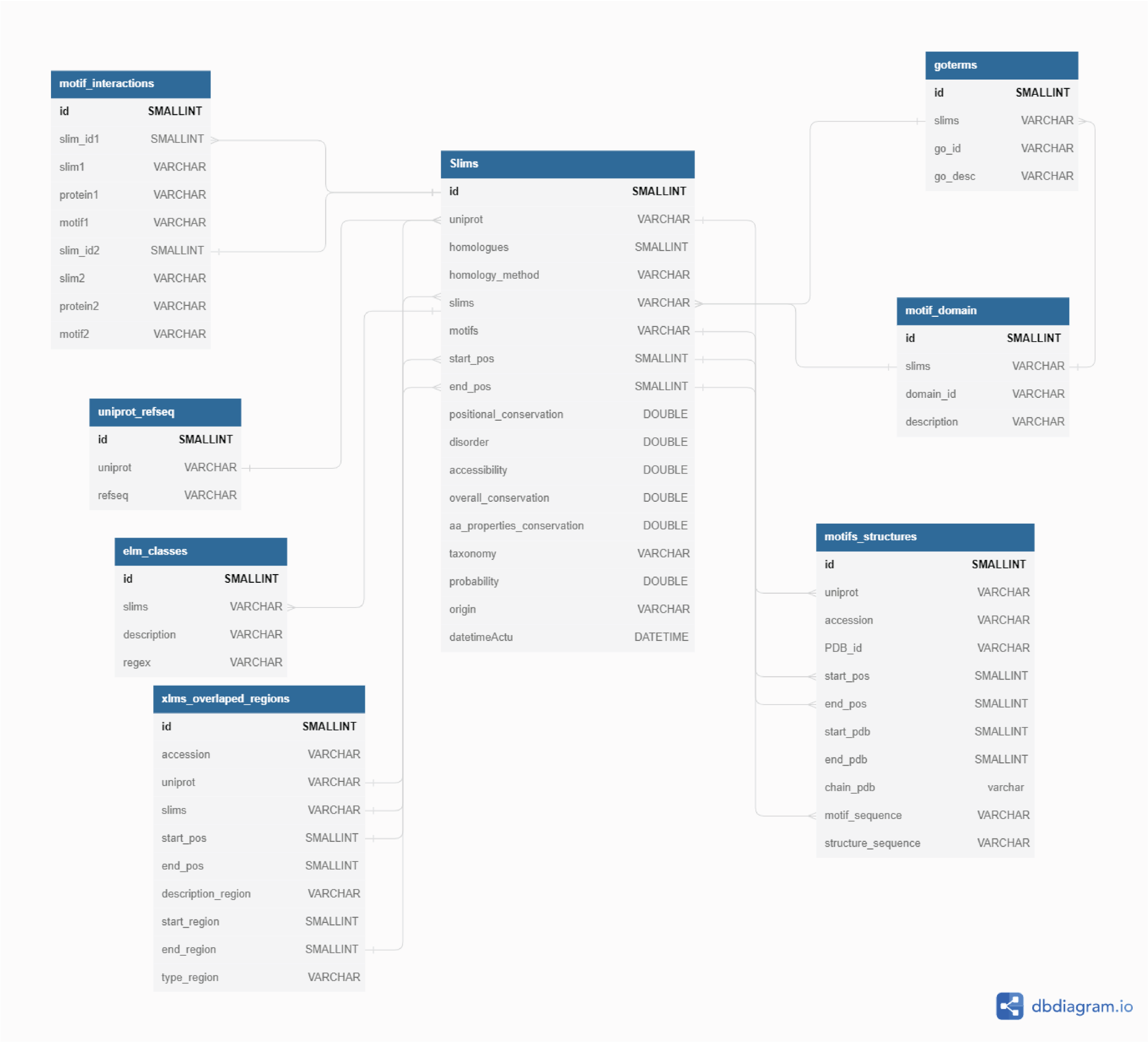


**Supplementary Figure S1: Table diagram of the evo-MOTiF database.**
Structure of the MySQL evo-MOTiF database. The table “Slims” contains results from the evo-MOTiF pipeline. “uniprot_refseq” is used to have correspondence between uniprot and refseq identifiers. “motif_domain”, “elm_classes” and “goterms” are extracted from ELM.eu.org to provide information about interactors, GO terms and descriptions of motifs from ELM.eu.org whereas “motif_interactions” is extracted from cross-linking associated with mass spectroscopy experiments from (Liu et al., 2017) to provide information about motif-motif interactions. “motifs_structures” contains results from SLiMs-SCAR, i.e. structural information about motifs. The figure was drawn using dbdiagram.io.

#### Supplementary Figure S2

##### evo-MOTiF database main motif page

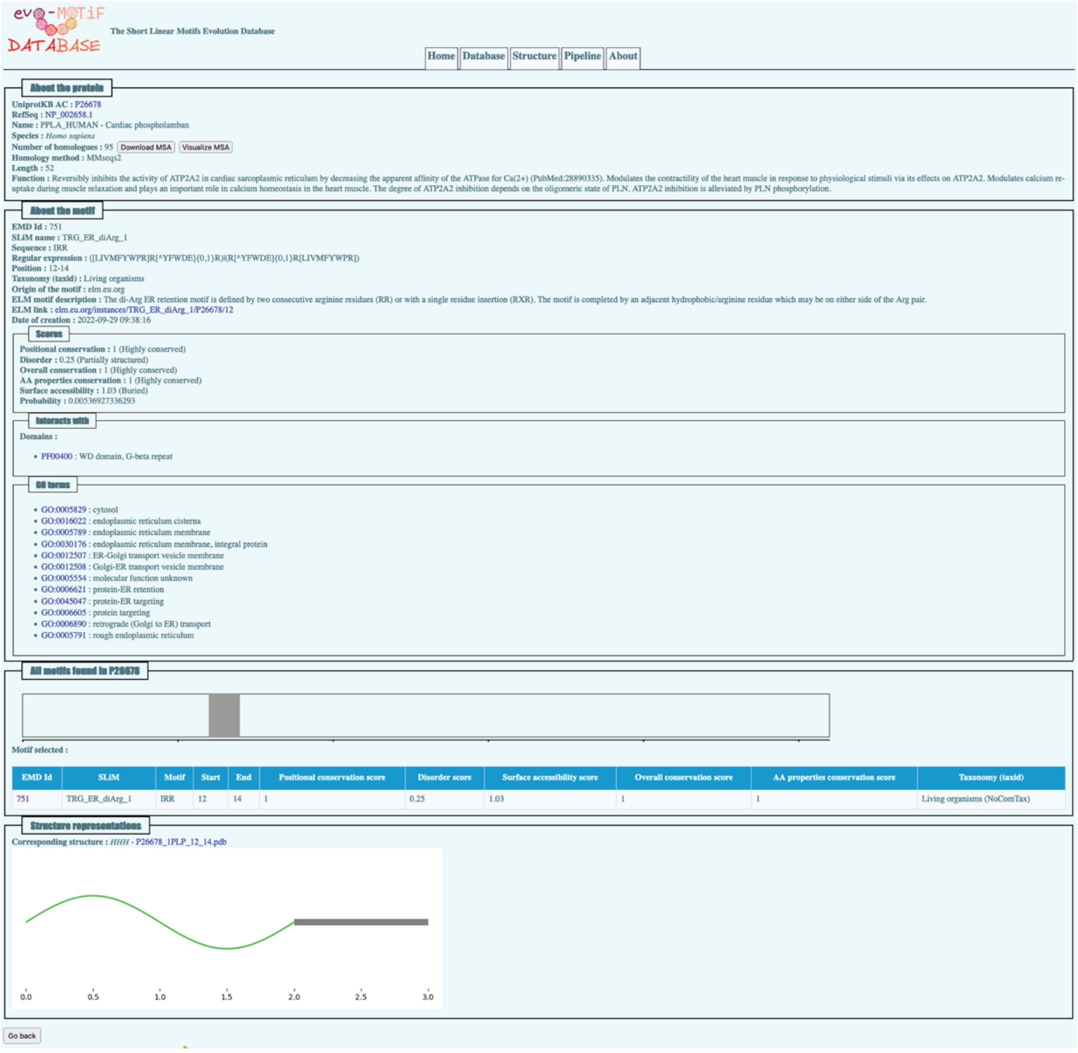


**Supplementary Figure S2: Main motif page of the evo-MOTiF database.** For each protein together with its known and experimentally verified SLiMs, a page exists in the evo-MOTiF database. At the top of the page, information on the protein itself is given, including its identifiers, the lengths, number of orthologs and which method was used to identify the orthologs. The multiple sequence alignment (MSA) of the protein family is accessible for download, as well as for live viewing in the browser (via the JSAV plugin, see Figure 3 of main text). This section is followed by information on the SLiM itself, which includes the name of the SLiM, the sequence and regular expression, the position in the sequence, the taxonomic identifier of the SLiM's host protein species, the number of orthologs, the SLiM is found in, the source of the SLiM, a description of its function, as well as a link to the source database. The next section shows the SLiM scores, including a classification of the different properties: these include order/disorder, conservation scores (amino acid property, positional, as well as overall conservation), surface accessibility, as well as the probability (derived from the ELM database). Information on the interacting domain (in case of ELM) or SLiM (in case of XL-MS data) is given, followed by Gene Ontology (GO) annotation of the SLiM (in case of ELM). In the lower part of the page, the protein sequence is schematically shown, with all the SLiMs it contains. The list of SLiMs is also given in tabular format below. Clicking on the SLiM icon or the database identifier will change to the main page of the SLiM in evo-MOTiF. At the very bottom, the structural representation of the SLiM is displayed and a link to the structure of the SLiM, as well as the protein containing the SLiM in colour, is given.

**Supplementary Figure S3**

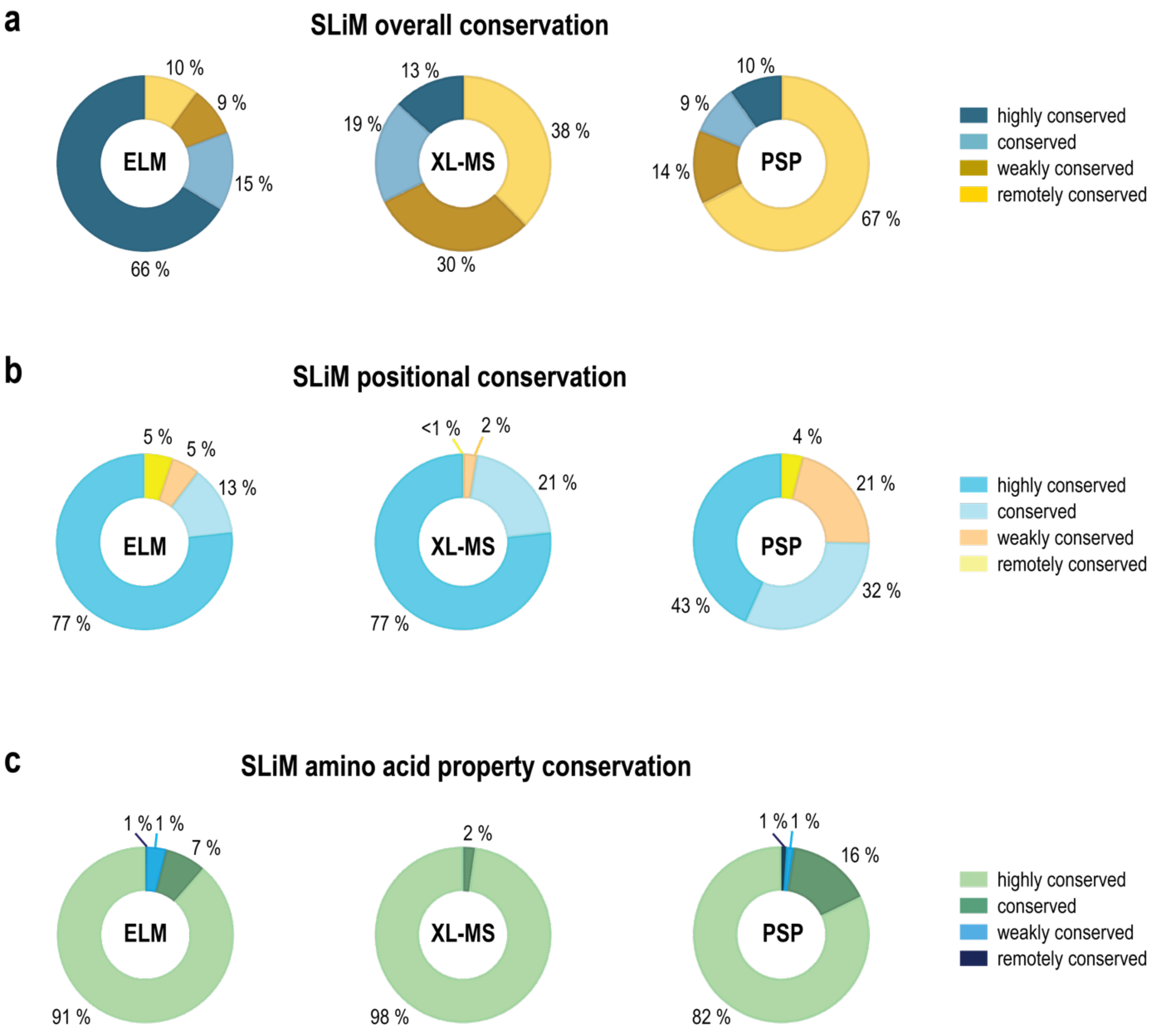


**Supplementary Figure S3: Overall (a) and positional (b) conservation scores for the SLiMs originating from the three different resources. (a)** Overall conservation of ELM, XL-MS and PhosphoSitePlus SLiMs. ELM contains a high percentage of highly conserved motifs (66%), while XL-MS and PhosphoSitePlus show more remotely or weakly conserved motifs. **(b)** Positional conservation of ELM, XL-MS and PhosphoSitePlus (PSP) SLiMs. The scores for positional conservation suggest generally higher conservation of ELM and XL-MS motifs (77%), while PhosphoSitePlus motifs display lower positional conservation. **(c)** SLiM amino acid property scores. Overall, the conservation of amino acid properties of SLiMs is high in all three resources.

**Supplementary Figure S4**

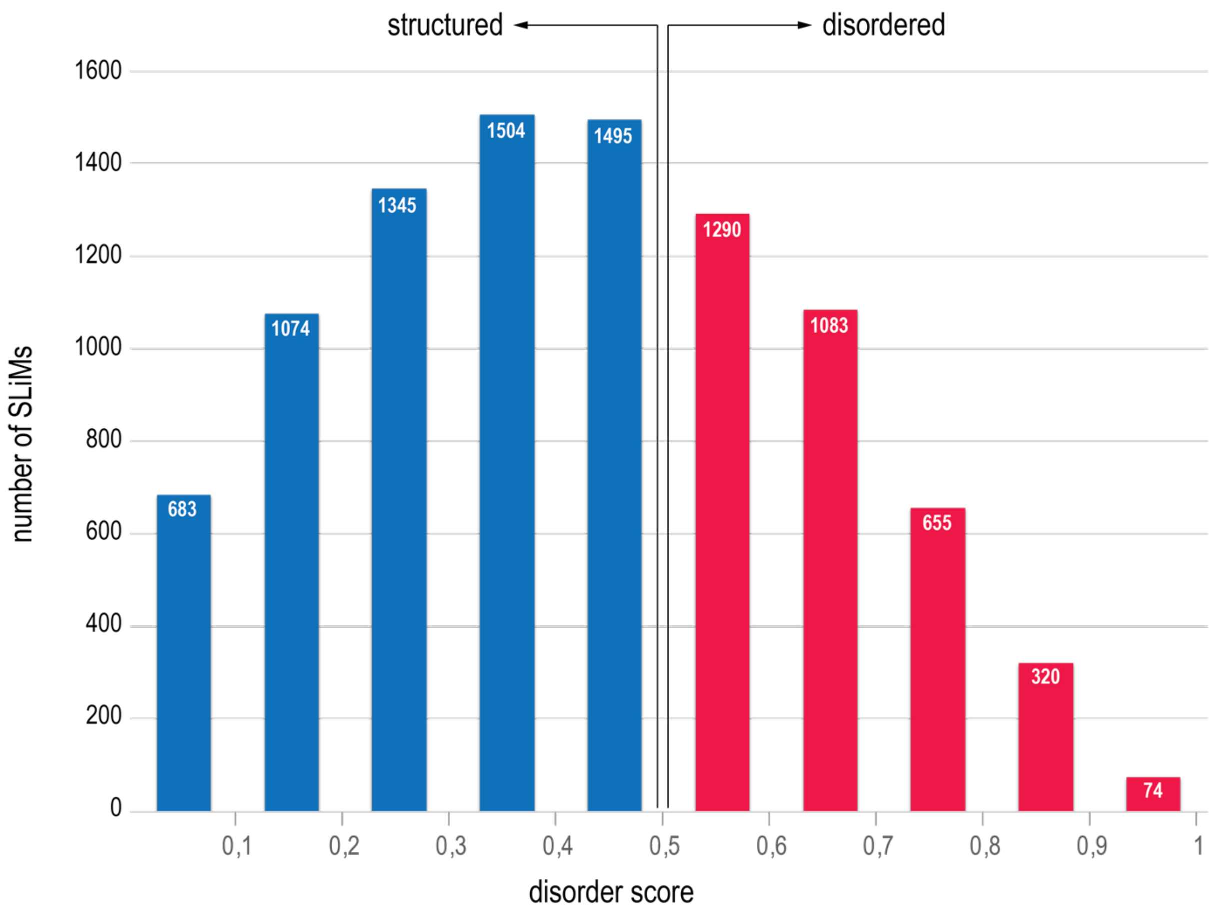


**Supplementary Figure S4: Frequency bar plot of structural SLiM features.** 60% of SLiMs from all three resources are categorised as 'structured' (blue bars). However, most SLiMs have a disorder score close to the disorder cut-off of 0.5.

**Supplementary Figure S5**

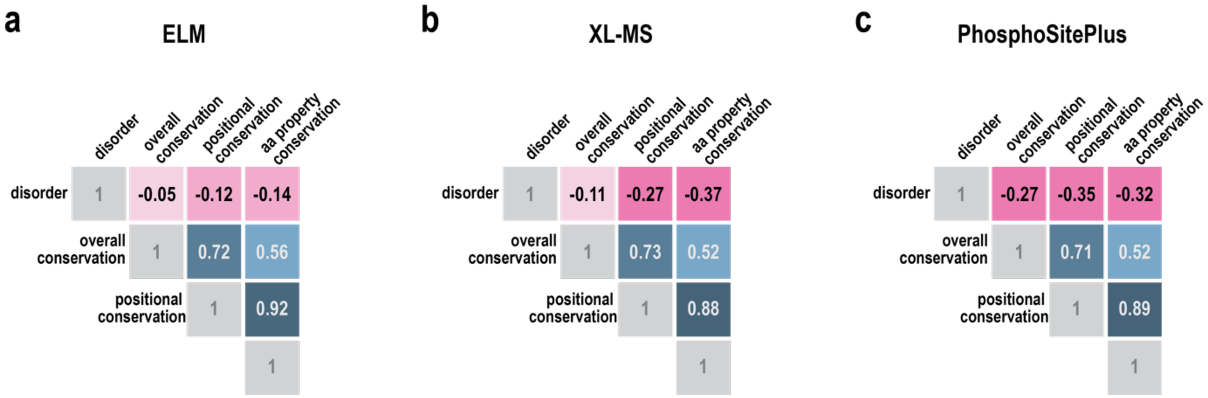


**Supplementary Figure S5: Correlation plots between disorder and conservation for all three resources. (a)** Correlation plot of ELM motifs. **(b)** Correlation plot of XL-MS motifs. **(c)** Correlation plot of PhosphoSitePlus motifs. The positive correlation between conservation scores is very high. There exists a negative correlation between disorder, as well as all three conservation scores. All correlation scores are statistically significant (p-value < 0.05; see also Supplementary Table S1 i for p-values of correlation scores).
